# Phase-dependent word perception emerges from region-specific sensitivity to the statistics of language

**DOI:** 10.1101/2023.04.17.537171

**Authors:** Sanne Ten Oever, Lorenzo Titone, Noémie te Rietmolen, Andrea E. Martin

## Abstract

Neural oscillations reflect fluctuations in excitability, which biases the percept of ambiguous sensory input. Why this bias occurs is still not fully understood. We hypothesized that neural populations representing likely events are more sensitive, and thereby become active on earlier oscillatory phases, when the ensemble itself is less excitable. Perception of ambiguous input presented during less-excitable phases should therefore be biased towards frequent or predictable stimuli that have lower activation thresholds. Here, we show with computational modelling, psychophysics, and magnetoencephalography such a frequency bias in spoken word recognition; a computational model matched the double dissociation found with MEG, where the phase of oscillations in the superior temporal gyrus (STG) and medial temporal gyrus (MTG) biased word-identification behavior based on phoneme and lexical frequencies, respectively. These results demonstrate that oscillations provide a temporal ordering of neural activity based on the sensitivity of separable neural populations.

## Introduction

Oscillations, or population rhythmic activity, reflect the waxing and waning of neural excitability such that individual neurons are modulated by an oscillation are primarily active on high excitable phases^1–3^. Previous studies have directly linked this phase-dependent neural activity to behavioral performance showing that target detection^4–7^ and accuracy in categorization tasks^8, 9^ are modulated by oscillatory phase. Besides accuracy, a few studies have also shown that oscillatory phase can modulate the categorization of ambiguous stimuli by biasing participants’ percept to one or another category based on the phase of presentation^10–12^. Improved behavioral performance has often been attributed to increased processing efficiency at oscillatory phases at which neural activity is increased^3, 13^. However, phase-dependent categorization biases cannot be explained by overall increases in activity (or increased processing efficiency) on specific oscillatory phases because increases in overall activity should not bias processing to one specific perceptual interpretation. Thus, it is unclear what neural mechanism underlies phase-dependent categorization.

Even though oscillations modulate neural excitability, not all neurons influenced by an oscillation reach activation exactly at the same time or phase. In fact, the phase-of-firing of a neuron is determined by an interaction between excitability changes due to oscillations and the neural sensitivity of a neuron to incoming signals^14^. Neurons that are sensitive to incoming signals will respond strongly and will therefore reach activation already at relatively low excitable oscillatory phases^1, 14^. In contrast, neurons less sensitive to the input will reach activation only at later, more excitable, phases. Neuronal sensitivity can be modulated by neuro-plastic changes induced through associative and statistical learning^15^. For example, neural populations representing more likely events in the world have higher sensitivity than populations representing less likely events^15, 16^. If this is true, populations representing probable events should be active at earlier, less excitable, oscillatory phases compared to populations representing less likely events which in turn could lead to phase-dependent categorization^17^.

Previously, we have shown that oscillatory phase in the theta frequency range can bias the categorization of an ambiguous syllable^11^. In that study, we presented an ambiguous syllable that Dutch participants could interpret as /dɑ/ or /xɑ/ (notation according to the international phonetic alphabet [IPA]). Originally, this phase-dependent categorization bias was attributed to an articulatory visual-to-auditory temporal difference between the two syllables (the visual-to-auditory articulatory delay of /dɑ/ is shorter than /xɑ/)^11, 18^. However, in Dutch, /d/ also has a higher frequency than /x/^19^, that is, /d/ is more probable than /x/. It is therefore possible that the categorization effect in this study was instead (or additionally) caused by an interaction between ongoing oscillations and the sensitivity of neural populations for these consonants^17^. Up to now, no study has investigated the role of event probability for the outcome of phase-dependent behavior. If oscillations provide a temporal ordering based on neural sensitivity^14, 17^, then one would expect phasic categorization effects to occur when there is a difference in event probability for two possible interpretations of an ambiguous stimulus. If this is so, we should view oscillations not merely as a gating operation opening and closing lines of neural communication^3, 13^, but rather as a rich source of representational space^20, 21^.

To investigate the relation between phase-dependent categorization and event probability we presented participants with words that varied in consonant, vowel, and word frequency. These were the four Dutch words *dat*, *gat*, *daad*, and *gaat* (see table 1 for translations, IPA notation, and event probability/frequencies). In this way, we manipulated event probabilities at different levels of analysis based on the recurrence of phonological and lexical elements in a language. By using computational modelling, psychophysics, and MEG we could investigate the influence of event probability on behavioral and neural responses to ambiguous stimuli (Figure 1). Computational modelling showed that phase biases categorization when words had different probabilities of occurrence. Moreover, the bias was dependent on the event probability within the specific level of analysis (phoneme or word level). These outcomes were verified using psychophysics and MEG.

**Table 1.**
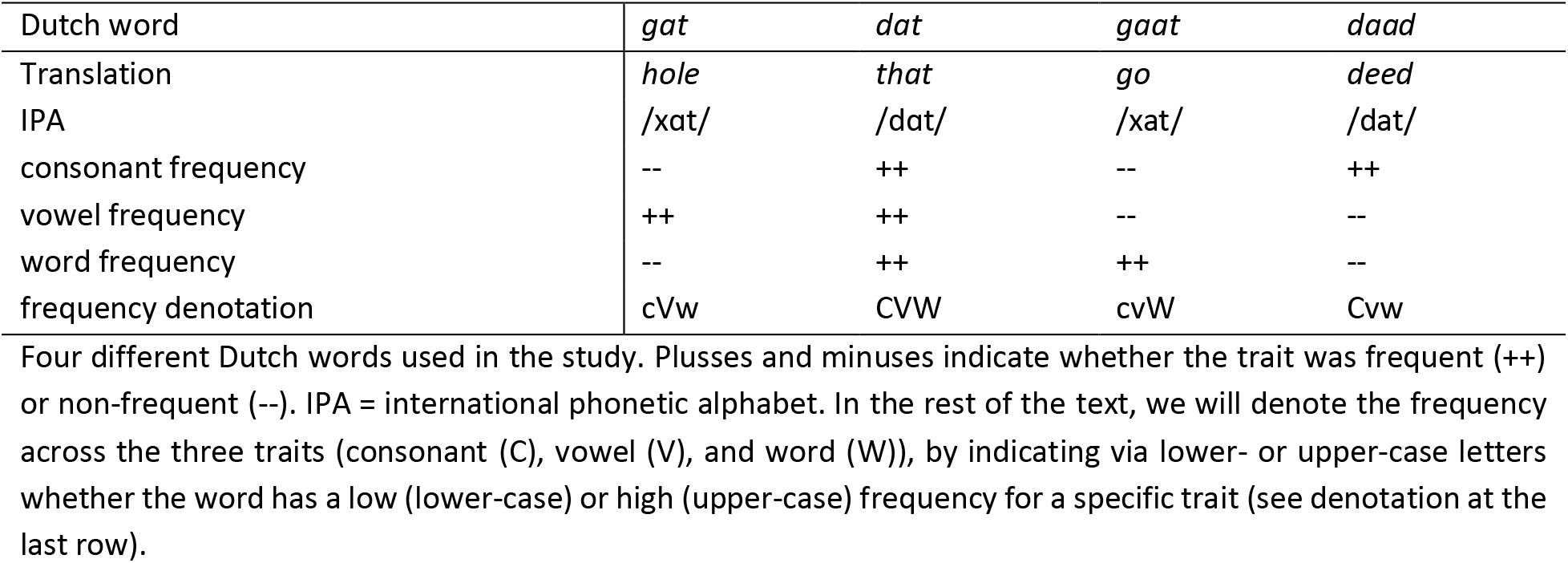
Stimulus materials

**Figure 1.**
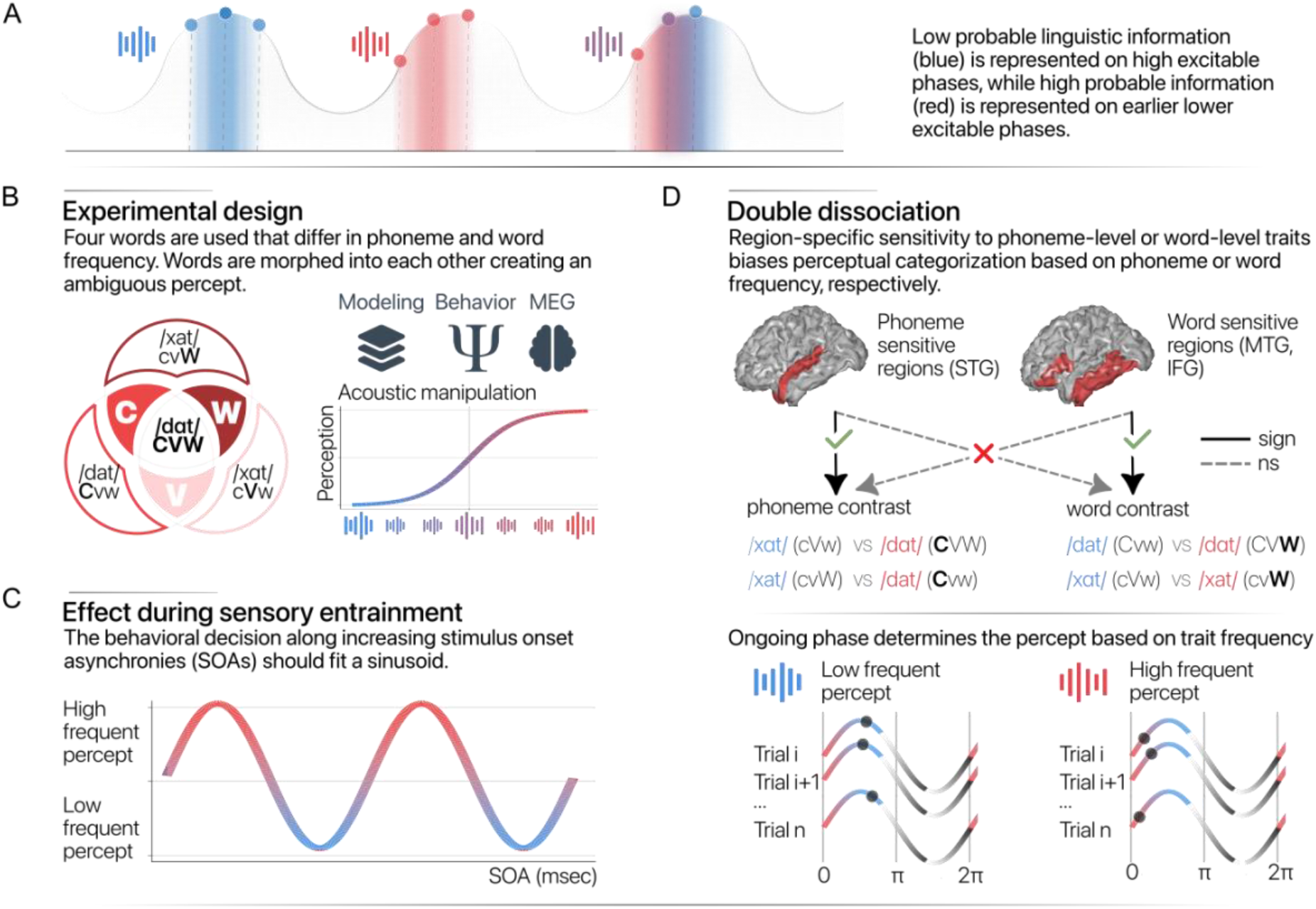
Overview of the current study. A) It is hypothesized that low probable linguistic information is represented at high excitable phases while high probable linguistic information is represented at low excitable phases. The perceived sound of ambiguous stimuli depends on the phase of presentation. B) Four words are used that differ in consonant (C), vowel (V), and word (W) frequency. Words are morphed into each other creating an ambiguous percept. C) Sensory entrainment locks neural rhythms to the rhythmic input and therefore the stimulus onset asynchrony (SOA) relative to the entrainment train should bias the percept to words containing low or high frequent linguistic information. D) In MEG a double dissociation is expected in which perceptual bias is governed by phoneme or word frequency for regions sensitive to phonemic or word features, respectively. Sign = significant; ns = not significant. STG = superior temporal gyrus; MTG = middle temporal gyrus; IFG = inferior frontal gyrus.

## Results

In the current study, we investigated whether combining oscillations with dissimilar ‘event’ probabilities – here phoneme and lexical frequency - can lead to phase-dependent categorization. First, we designed a computational model whose goal was to categorize incoming sensory input. To directly link the outcome of the computational model to empirical findings and quantify phase-dependent categorization, we presented the model with the stimuli used in the psychophysics and MEG experiment. In the psychophysics study, we presented an entrainment stream at 6.25 Hz after which an ambiguous word was presented at a variable stimulus onset asynchronies (SOAs). Assuming an entrained oscillation, these SOAs match to ongoing oscillatory phases^22^. We generated ambiguous words by creating 10 equally spaced morph levels along each dimension (that is varying either the consonant /x/-/d/ or the vowel /a/-/ɑ/). This procedure resulted in four different morph scales: /xat/-/dat/, /xɑt/-/dɑt/, /dɑt/-/dat/, and /xɑt/-/xat/. Participants categorized each morphed stimulus as one of either word. The most ambiguous morph was defined individually for each participant by fitting a psychometric curve to their responses and selecting the morph closest to 50% categorization. In the MEG experiment, stimuli were not preceded with an entrainment train, but were presented at random SOAs.

## Computational model

The computational model was an extension of the Speech Tracking in a Model Constrained Oscillatory Network (STiMCON) model introduced in ^17^. This model integrates temporal tracking together with the tracking of the content of speech using both temporal and content information. Input activation levels at a given time (A*_l,T_*) are governed by the following function:

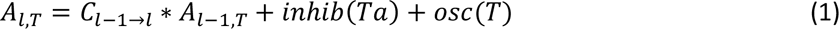

in which C represents the connectivity patterns between different hierarchical levels (l), T the time in milliseconds, and Ta a vector representing the times of individual nodes within the inhibition function (see online methods). Input activation is thus determined by activations from lower levels as well as an inhibition and an oscillation function. Individual Ta node values are set to zero as soon as activation of a node reaches activation threshold (default threshold = 1). This function first ensures non-linear supra-threshold activation after which the node is temporally inhibited.

In the current implementation, we introduced two different levels of analysis: a phoneme and a word level. Both levels receive input from the input level but have their own connectivity with the input and their own node sensitivity. The input is modelled as the individual words: /xat/, /xαt/, /dat/, /dαt/, and an empty word node is used for the entrainment train. For the phoneme level, we represented the phonemes /x/, /d/, /a/, /α/, and an empty phoneme node. Connectivity for the input-to-phoneme level was set to one when the phoneme was part of the input word (the entrainment stimulus was connected with a one to the empty phoneme node). The input-to-word level connectivity consists of an identity matrix (each word loads with one on the word level). Sensitivity of individual nodes to input was varied by reducing the activation threshold for the more frequent phonemes (/d/ and /α/) and words (/dαt/ and /gat/) in their respective analysis level (activation thresholds were parametrically reduced between 0.01 - 0.1). In all simulations, we extracted the categorization response of the model by determining the deciding node that was active first after stimulus presentation. For the two categorization options, we coded one node as a 0 and the other node as a 1 (if both nodes were active simultaneously or no node was activated at all we assumed that the model would guess and set the value to 0.5 for the psychophysics simulation or choose randomly 0 or 1 for the MEG simulation). For the psychophysics experiment, we assumed that the output of the model reflects the average outcome of the phoneme and word level.

We let the model run through the psychophysics and MEG experiments. In both experiments, all morphs are initially presented at random moments to generate a psychometric curve and to determine the most ambiguous stimulus that will be used for the main experiment (see online methods for more details). To imitate this procedure, input was presented at different amplitude proportions of two words (e.g. for /xat/-/dat/ morph with 90% of /dat/, /xat/ was presented with an amplitude of 0.1 and /dat/ at 0.9) evenly distributed across all phase values. For each morph, we averaged the node responses of the model across the repetitions of the same morph (and across the two levels). For all word morph spectra, we could reliably fit a psychometric function and extract the most ambiguous morph (Figure 2A). For the second part of the psychophysics experiment, the model was presented with an entrainment train of empty words after which we presented the most ambiguous morph stimulus at different SOAs. Results show that only for morph spectra in which the two morphed traits had opposing event probabilities or frequency (i.e. /dɑt/ and /xɑt/ [CVW vs cVw] and /dɑt/ and /dat/ [CVW vs Cvw]) a phase-dependent categorization performance developed across all sensitivities tested (Figure 2B and 2C).

**Figure 2.**
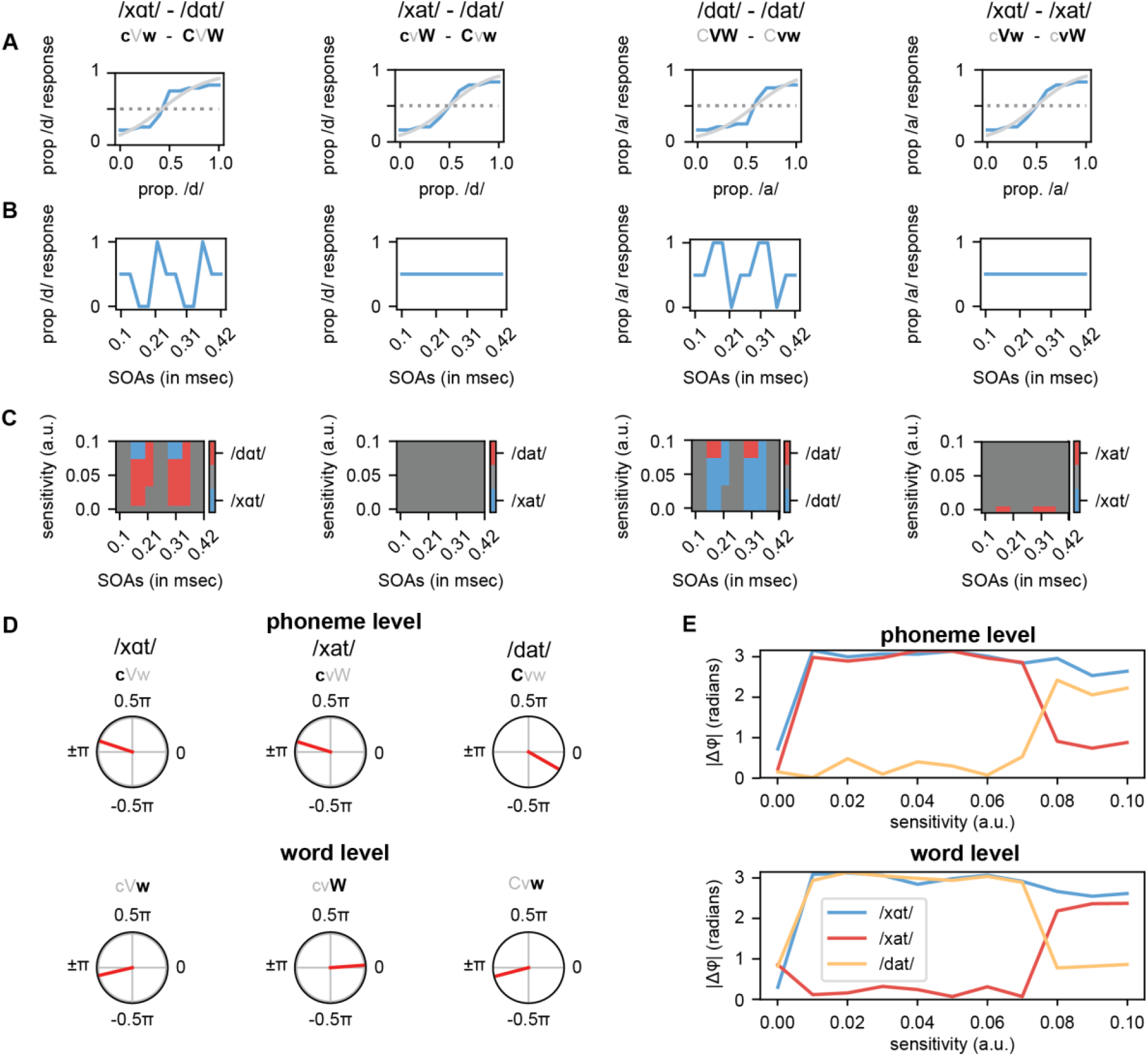
Outcome of the computational model. A) Psychometric functions for the four different morph dimensions (sensitivity = 0.09). Blue lines represent the model output; grey lines show the psychometric fit. B) Response of the model to the most ambiguous morph of A presented at different stimulus onset asynchronies. C) Response choice of the model during entrainment (as in B) for different sensitivity levels. D) Phase difference between the average phase of the three different response choices and /dɑt/ (sensitivity = 0.07). E) Phase differences (as in D) for different sensitivity levels.

For the MEG experiment, phase does not have to be inferred from an entrainment train, but rather can be estimated from the recorded regions directly. To simulate this experiment, ambiguous morphs were presented to the model at random phases (repeated for 1000 repetitions). Only the consonant morphs along the /dɑt/-/xɑt/ and /dat/-/xat/ spectra were used in the MEG experiment. For each of the two ambiguous morphs, the phase was extracted together with the categorization response based on the node activation of the phoneme or word level. For the main MEG experiment, we are not hypothesizing about the absolute phase, as we have no hypothesis about the exact phase (see^6, 11^), but rather in *relative* phase difference between a more-likely versus less-likely event. Therefore, we took the phase difference between the word /dɑt/ which has a high frequency on all trait dimensions (CVW) and the phase of the other response options. For the phoneme level, the model showed high phase differences of around π when the ambiguous morph was interpreted as either /xat/ or /xɑt/, but low phase differences of around 0 when the model interpreted the morph as /dat/ (Figure 2D). In contrast, for the word level, we found π phase differences for the categorization choices /xɑt/ and /dat/, and 0 phase difference for the choice /xat/. Thus, phase differences were low when both words had high frequency traits within the level of analysis (high frequency phonemes in the phoneme level [/dɑt/, CVW vs /dat/, Cvw] and high frequency words in the word level [/dɑt/, CVW vs /xat/, cvW]). Phase differences were high when the words had different frequency traits (high versus low frequency phonemes in the phoneme level [/dɑt/, CVW vs /xɑt/ [cVw] and /xat/, [cvW] and high versus low frequency words in the word level [/dɑt/, CVW vs /xɑt/ [cVW] and /dat/, [Cvw]). In MEG, this pattern of results could correspond to phase differences in different neural sources that analyze phoneme- and word-level representations, respectively. These phase differences were more pronounced when the sensitivity level changes were relatively low (Figure 2E).

## Psychophysics experiment

We conducted a consonant and a vowel version of the experiment by morphing either the consonants of the words or the vowels of the words, creating four morphs: /xat/-/dat/, /xɑt/-/dɑt/, /dɑt/-/dat/, and /xɑt/-/xat/. During the first part of the experiment, we presented stimuli across all morphs to create an individual psychometric curve along the consonant (Figure 3A) and vowel dimension (Figure 3D). Only participants for which we could reliably extract an ambiguous stimulus via fitting a psychometric curve could participate in the main experiment.

**Figure 3.**
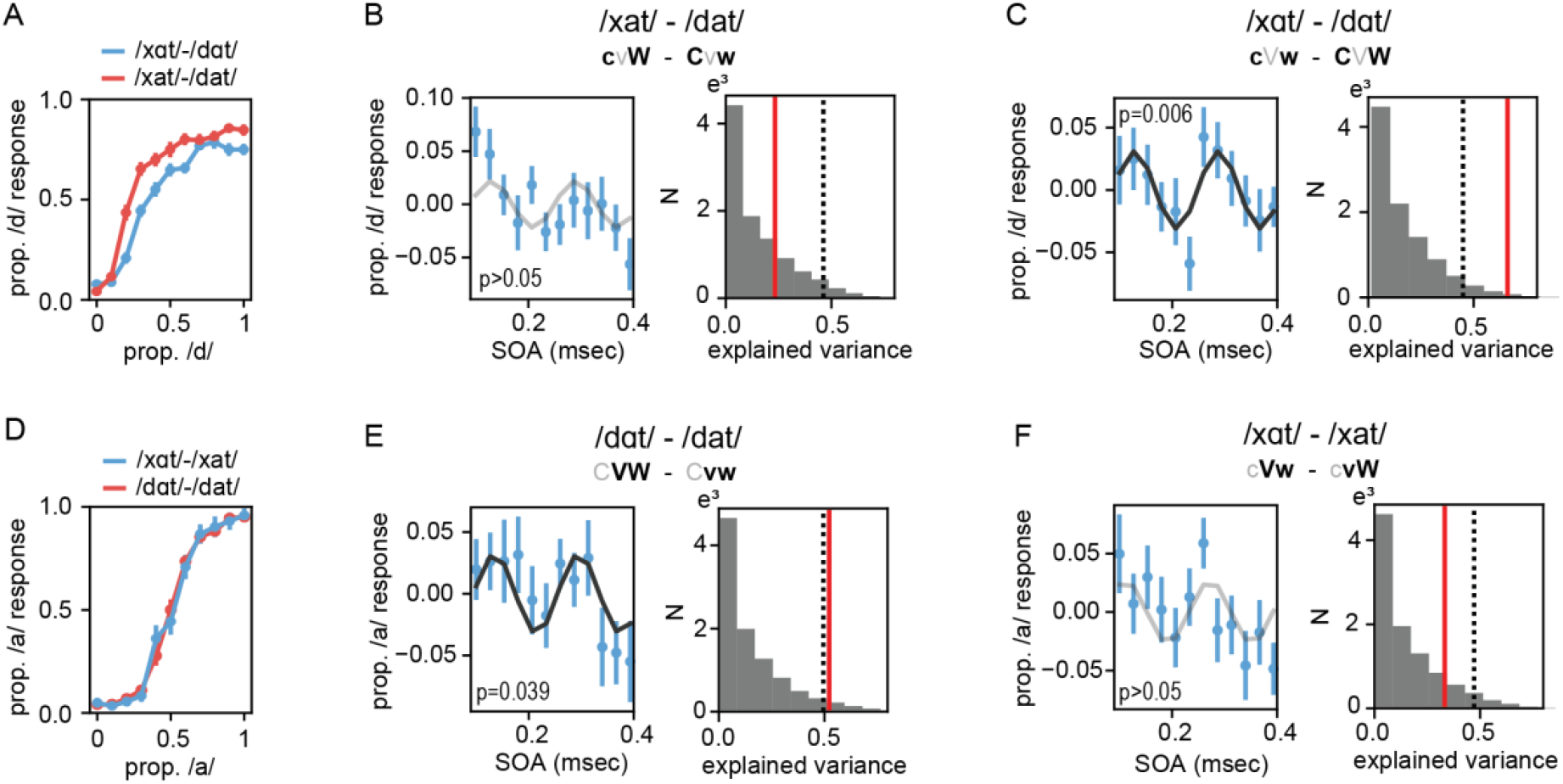
Behavioral results. A) Psychometric curves for the consonant experiment (separate lines for the two vowel types). B) Outcome of the main experiment for the /xat/ - /dat/ spectrum. The left panel shows the average demeaned time course across stimulus onset asynchronies (SOA). The right panel shows a histogram of the null distribution together with the observed explained variance of the sinusoid fit (red line) and the 95^th^ percentile (dotted line) of the null distribution. C) Same as B for the /xɑt/-/dɑt/ spectrum. D-F) Same as A-C for the vowel experiment. Error bars indicate the standard error of the mean. Black lines indicate the best fitted curve (gray if not significant).

In the main experiment, participants listened to rhythmic broadband noise bursts presented at 6.25 Hz after which an ambiguous word was presented. The SOAs at which ambiguous words were presented ranged between 0.1 and 0.4 seconds in 12 equidistant steps (spanning exactly two cycles of 6.25 Hz). Participants had to indicate which word they heard. For the consonant experiment, we found that a 6.25 Hz sinusoid fitted to the data yielded a higher explained variance than expected by chance for the /xɑt/-/dɑt/ morph (p = 0.006, r^2^ = 0.661; Figure 3C), but not for the /xat/-/dat/ morph (p > 0.05; Figure 3B). For the vowel experiment, we could significantly fit a sinusoid for the /dɑt/-/dat/ (p = 0.039, r^2^ = 0.523; Figure 3E), but not the /xɑt/-/xat/ morph (p > 0.05; Figure 3F). In sum, we could only fit a significant curve for morphs in which both varied traits had opposing frequencies in the word pairs, that is for the /xɑt/ - /dɑt/, cVw – CVW, morph and the /dɑt/ - /dat/, CVW – Cvw, morph. This was in line with the outcome of the computational model (Figure 2).

## MEG-experiment

The output of our computational model showed that event probability determines whether we can expect phase-dependent categorization. Moreover, the phase-dependent categorization was different for the phoneme and the word level of analysis. However, in the psychophysics experiment, all responses are based on an integration across both levels of analyses as there is only one behavioral output. To bridge this gap, we designed a MEG study in which the phase-dependent categorization effects at different levels of analysis could be source localized to cortical regions that are known to correspond to varying levels of linguistic computations^23, 24^. Indeed, earlier auditory regions such as superior temporal gyrus (STG) are more sensitive to phoneme content, such as vowel and consonant traits, while regions higher in the auditory hierarchy, such as medial temporal gyrus (MTG) and inferior frontal gyrus (IFG) are sensitive to lexical representation and temporal integration, respectively^23, 25–28^. If phoneme or word frequency relate to phase-dependent categorization, the phase of ongoing oscillations in distinct cortical regions should bias categorization based on the level of analysis of that region. To test this hypothesis, we presented the ambiguous morphs of the consonant experiment to Dutch participants while recording their neural activity with MEG and source localizing the response to the STG, MTG and IFG.

First we looked at the overall power response in the pre-stimulus period (see Supplementary Figure 1 for post-stimulus responses). All analyses were based on using an array-gain beamformer which corrects for center-of-head biases without the need of a baseline^29, 30^. To limit computational resources, we focused on the first PCA of all grid points part of the ROI quantifying the component explaining the most variance in the ongoing data. In all ROIs we found theta peak (peak values: 8.4 Hz, 8.6 Hz, and 8.2 Hz for STS, MTG, and IFG, respectively) across the whole pre-stimulus window, but this peak was weaker for the IFG (Figure 4). We compared pre-stimulus power values dependent on the response of the participant for the ambiguous stimuli but found no differences (all p > 0.573).

**Figure 4.**
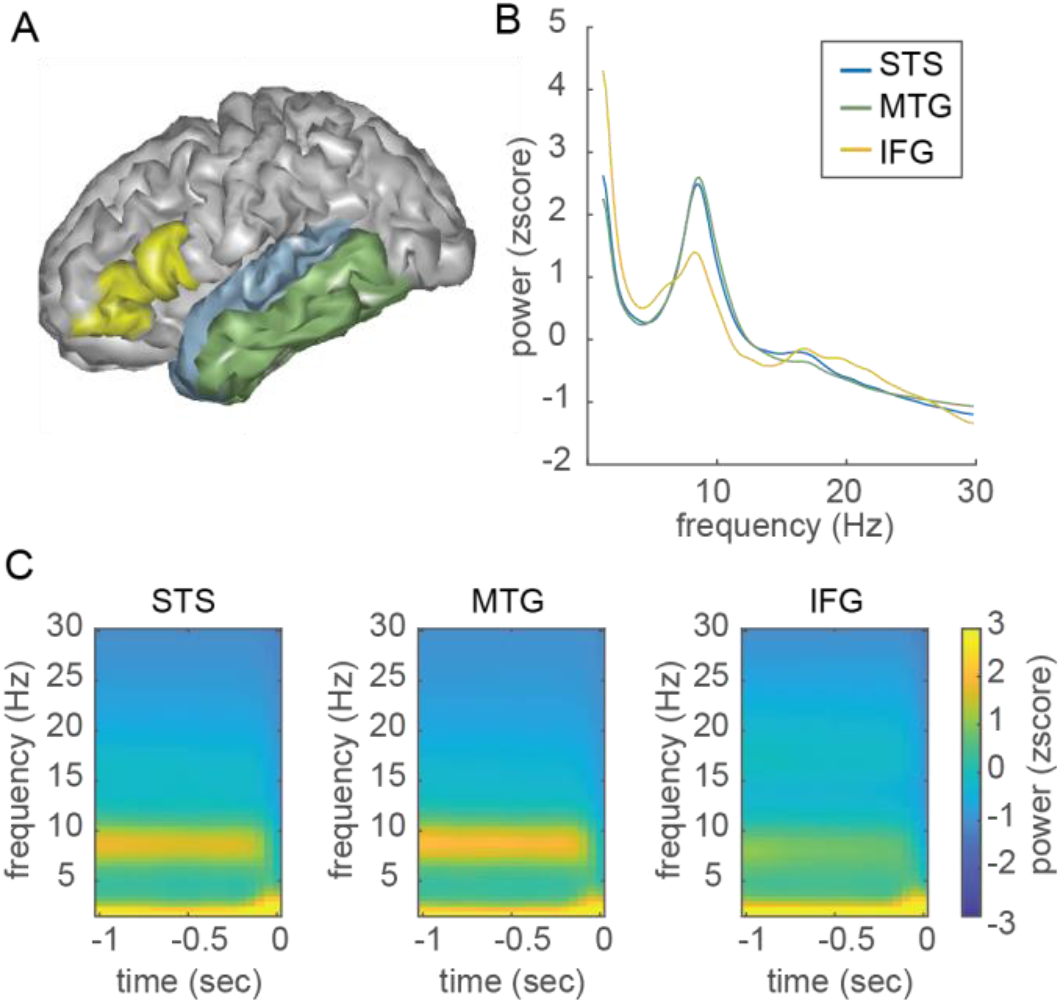
Pre-stimulus power in regions of interest. A) The three regions of interest. B) Power spectra averaged for the - 1-0 sec time window averaged across the two ambiguous stimuli and response choices. C) Time-frequency response averaged across the ambiguous stimuli and response options. Note that data is padded from time point 0 on (explaining the sharp power drop around zero).

For all participants, we could individually determine the most ambiguous morph in the first part of the experiment and all, but one could maintain an ambiguous percept throughout the experiment (Figure 5A-B). We investigated whether phase-dependent categorization was determined by phoneme or word frequency for each ROI separately. At each pre-stimulus time-frequency point, we performed a circular-linear correlation between the pre-stimulus phase and response type (low or high trait frequency; that is, for the consonant contrast /xɑt/ [cVw] and /xat/ [cvW] vs /dɑt/ [CVW] and /dat/ [Cvw]; for the word contrast /xɑt/ [cVw] and /dat/ [Cvw] vs /xat/ [cvW] and /dɑt/ [CVW]). To correct for multiple comparisons, we ran cluster-based statistics^31^. For the consonant contrast we found a significant effect of consonant frequency in the STG (Figure 5C; cluster statistic: 69.655; p-value: 0.022; frequency range: 4.4 - 9.0 Hz; time range: -0.30 - -0.10 sec; peak t(21)-value: 4.119 at 7.617 Hz, -0.20 sec), but not in the MTG or IFG (p > 0.05). For the word contrast, we found a significant effect of word frequency in the MTG (Figure 5C; cluster statistic: 66.138, p-value: 0.019; frequency range: 4.4 - 8.5 Hz; time range: -0.25 – 0 sec; peak t(21)-value: 3.589 at 6.445 Hz, -0.10 sec), but not in the STG or IFG (p > 0.05). In sum, we found a double dissociation between ROI and trait type.

**Figure 5.**
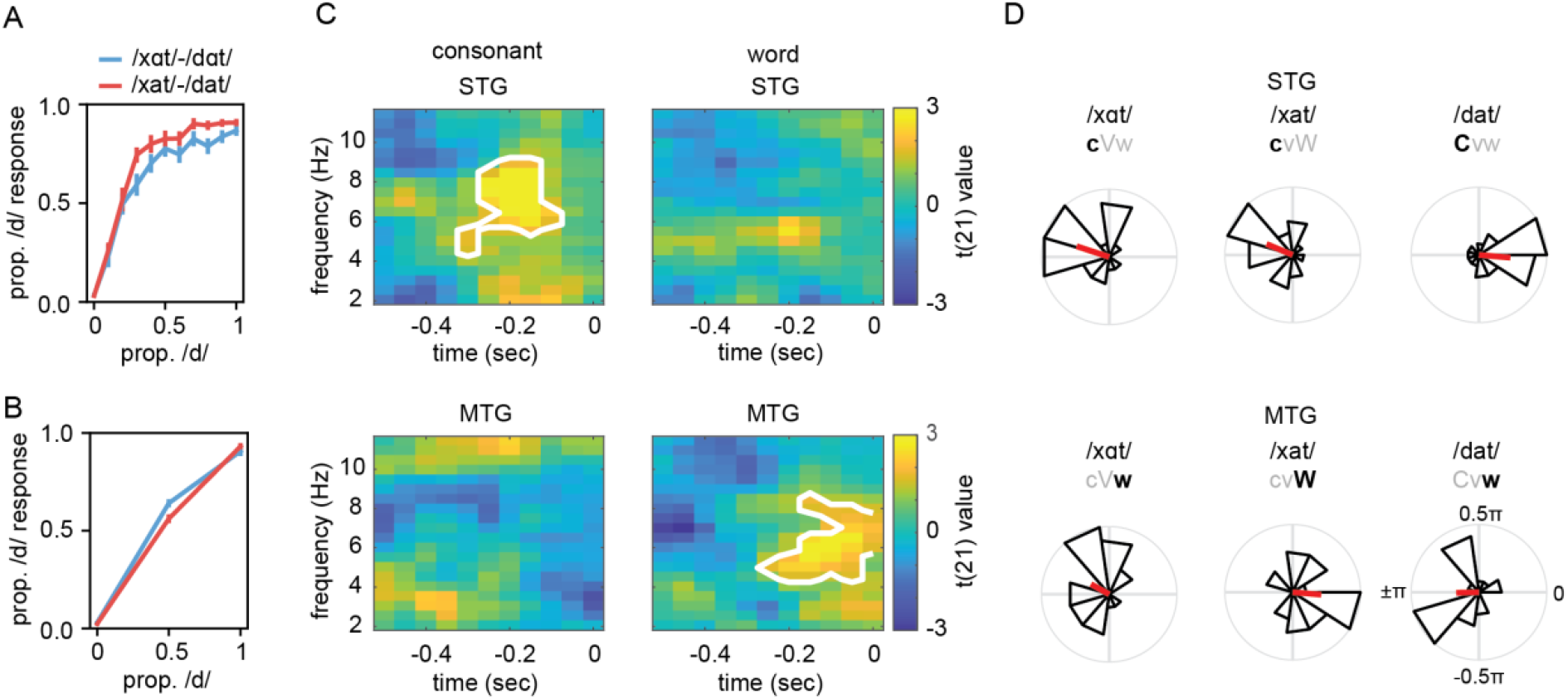
MEG results. A) Behavioral responses in the MEG experiment for part one of the experiment. B) Behavioral response for the main experiment. Error bars reflect the standard error of the mean. C) Circular-linear correlation results show that in STG phase determines the percept of consonant frequency (top) while in MTG phase determines the percept of word frequency (bottom). White outline indicates significance (p < 0.05) according to the cluster-based statistical test. D) Phase difference with CVW /dɑt/ at the individual peak time-frequency point in C. In STG, words with high frequency consonants (Cxx words) had a phase difference of 0, while words with low frequency consonants (cxx words) had a phase difference of around π (top). In MTG, high frequency words (xxW words) had a phase difference of 0, while low frequency words (xxw words) had a phase difference of around π (bottom).

To further evaluate the exact phase differences for each ambiguous sound, we computed the average phase at which participants heard either of the two words. Phases were extracted for each individual’s peak time-frequency point within the significant cluster (for both morphs separately). The exact MEG phase was not expected to be identical across participants as it is difficult to determine excitability levels of an oscillation from MEG phase and individual stimulus processing times might differ. Rather, high event probabilities should be represented at the same phase, while low event probabilities are represented at the opposite phase within each participant. We took the word /dɑt/, which had high frequencies on all traits (CVW word), as a reference word and took the phase difference between the average phase at which participants heard /dɑt/ and one of the three words (ambiguous words perceived as /xɑt/, /xat/, or /dat/). We expected that for high frequency traits the phase difference would be zero, while for low frequency traits the phase difference would be around π relative to the reference word. To test this, we performed a v-test that tests for non-uniformity with a specific direction for each of the three words per ROI. To generate a p-value that combines the three values (as we expected all three contrasts to be significant), we multiplied the three probabilities yielding the probability of all three events happening at the same time (assuming independent tests).

In STG, we found that the average phase difference for the three words /xɑt/, /xat/, or /dat/ was 0.90π, 0.87π, and -0.027π respectively. The three phase differences were close to the expected phase differences: the low frequency consonants having a phase difference of π, while the high frequency consonant having a phase difference of zero (Figure 5D). The individual tests showed significance (/xɑt/: vstat = 10.56, pval < 0.001; /xat/: vstat = 8.47, pval = 0.005; /dat/: vstat = 10.18, pval = 0.001 respectively) as well as the combined probability (p < 0.001). In MTG, the average phase difference for the three words /xɑt/, /xat/, or /dat/ was 0.84π, -0.026π, and -0.99π respectively. The phase differences were close to the expected phase differences: the low frequency words having a phase difference of π and the high frequency word having a phase difference of zero. The individual tests showed significance (/xɑt/: vstat = 5.91.56, pval = 0.037; /xat/: vstat = 9.26, pval = 0.003; /dat/: vstat = 7.36, pval = 0.013 respectively) as well as the combined probability (p < 0.001). Thus, also for the individual words, the phase differences matched the neural sensitivity of the underlying region.

## Discussion

In the current study we used computational modelling, psychophysics, and MEG recordings to demonstrate that variations in neural sensitivity to event probability operationalized as phoneme and word frequency can result in phase-dependent perceptual categorization. We showed that ambiguous words presented at different phases, either through neural entrainment or by extracting the phase from MEG, are interpreted as one or another word depending on the time or phase of presentation. Moreover, in the MEG data we could dissociate these effects to separate cortical regions: phase-dependent categorization in STG depended on phoneme frequency, while word frequency modulated phase-dependent responses in MTG. The behavioral findings and the double dissociation between STG and MTG responses matched the results from a computational model that uses oscillations, together with varying neural sensitivity, to capture categorization responses. These results demonstrate that the neural phase code relies on ordering based on neural sensitivity, directly linking phase-coding to behavioral outcomes in a categorization task.

Most studies investigating the direct link between ongoing oscillatory phase and behavior have focused on assessing the role of oscillatory phase in modulating overall performance measures, such as accuracy^8, 32^, detection^4, 5, 33^, or reaction times^34^. These studies are all based on the assumption that oscillations modulate overall firing rates and subsequent neural processing should be optimized at phases where neural excitability is high^3^. We here argue that this view might be too simplistic and does not provide the full picture of the role of oscillations for neural computation. Instead of merely providing windows of processing opportunity^3, 13^, oscillations provide a means to organize the complex neural dynamics by activating neural populations at different neural phases^14, 20, 35^. This organizational principle of phase coding has extensively been shown with invasive recordings in animals^36, 37^. Moreover, prior computational modelling has shown the computational benefit of this organizational principle, as it effectively increases the representational space in the brain^35^ and changes the formal expressive power of a system^38–40^.

It has been an open debate what are the organizational principles of phase coding, that is, what determines when a neuron is activated^37^. In working memory paradigms, sequence order has often been implied to be the main organizational principle of phase coding^41, 42^. This is based on studies primarily in rats that show phase precession in which the order of the phase code is linked to the order of upcoming locations in an explorative maze task^36^. In our study, no sequence order can be imposed. Nonetheless, we find that the behavioral outcome of a phase code is based on the frequency of phonemes and words in the Dutch language by using computational modeling, psychophysics, and MEG. This finding suggests that sequence order is not the only principle that can generate a phase code. Instead, overall event probability of words within a language modulates the neural sensitivity and can also impose a phase code^17^. While the present study is focused on event probability based on the overall frequency of information in a language, we hypothesize that this finding can be extended to event probability that also depends on contextual knowledge. Evidently, contextual event probability and sequential order are related: events that are going to happen earlier in the future are more probable in the short-term. Moreover, events occurring in the near future have a higher behavioral relevance. Both probability and behavioral relevance could have a consequence for how excitable individual neural populations are. Therefore, excitability shifts, rather than solely order or event probability, could be the core principle that organizes the phase code.

The brain’s sensitivity to varying levels of event likelihood based on the statistics in the world has been shown in a plethora of studies which demonstrate that the brain is more sensitive to stimuli that are more likely^16^. However, the consequence of a probability manipulation in combination with oscillatory dynamics has rarely been studied. We here provide the first behavioral evidence showing that event probability and oscillations together provide a phase code which activates event representations on different phases based on their likelihood. It is unknown how strongly event statistics modulate neural sensitivity. This could potentially be relevant for the behavior of a neural system as our computational model suggests a non-linear change in phase-dependent perceptual outcomes for increasing neural sensitivities (Figure 2E). Probability modulations as tested in the current study are rather static and depend on word probability in language, which has been learned over the course of one’s life. It would be interesting to also investigate whether these effects are dynamic by varying event probabilities within the course of an experiment. If this manipulation also leads to similar phase-dependent categorization, phase codes would not only be adjusted solely based on long-term hard-coded changes in excitability, but also based on dynamically changing excitability levels that rely on top-down feedback^17^.

It has been proposed that speech comprehension involves first parsing speech into separate temporal chunks using neural oscillations^43–45^. This segmentation is hypothetically done by aligning theta band oscillations with syllables in an ongoing speech stream^44^. In this way, one can parse and identify individual syllables and use them for higher order linguistic operations^46^. In our study, it is difficult to separate segmentation or ‘chunking’ from any kind of process of interpretation. If a word is first ‘segmented’ or ‘chunked’ by theta oscillations, the information about the phase would be lost in subsequent operations. However, in our study, it is exactly the phase of the theta oscillation that determines how a word is interpreted. Note also that the reported effects are strikingly close to the 6.25 Hz frequency we found in our previous study^11^. Therefore, our study shows that segmentation or chunking through oscillations cannot be treated as a wholly separate process from word recognition, because oscillations also provide a categorization mechanism alongside any potential segmentation or ‘chunking’ operation (see also ^47,48|49^).

We have previously argued that during natural speech and language processing temporal information can be used to infer information content, in other words, time can be a cue for content (also see ^17, 50^). This time-content relation is governed by the observation that words that are more likely in the current context are uttered with shorter inter-word-intervals^17, 51, 52^. Combining this observation with theories of oscillatory tracking results in more likely words being naturally presented at earlier, less excitable phases, which we confirmed with our computational model^17^. This type of phase code can aid speech comprehension: when information is ambiguous, the phase of an oscillation, and thereby the time of word presentation, can be used to determine the percept. Notably, our current study does put some limitations on this use of phase-dependent categorization. Our behavioral analysis shows that this phase-dependent categorization works mostly when trait features across the word are all either frequent or non-frequent. Yet, it is not clear how likelihood information that varies based on the level of linguistic analysis interacts with timing in natural speech. To investigate this, one would need to show how top-down feedback modulates phase-dependent categorization in sentence context as top-down feedback strongly influences the sensitivity of individual word nodes as well as the timing of speech^17^.

In our computational model, we postulated a separate phonological and lexical level of analysis. This is in line with neuroscientific results, which show that speech analysis is split up in separate analytical levels across the cortex^23, 24, 53^. Specifically, it is suggested that STG is sensitive to phonological content, while MTG is sensitive to lexical access and word content. It is known that anatomical and functional connectivity between these regions operate via early auditory cortex. There are direct connections between early auditory cortex and STG and MTG, but also connections between STG and MTG^54–56^. Our model currently relies primarily on the direct connections. An additional area strongly involved in the language network is the IFG. We could not find any phase-dependent categorization in the IFG. This null finding is not necessarily surprising as IFG has mostly been associated with higher-order language processes that involve temporal integration across words and syntactic analysis^27, 28, 57^. IFG might simply be less sensitive to phonological or word-level likelihood differences. Alternatively, it is possible that we did not find phase-dependent effects simply because there is too much temporal variability in the IFG responses to speech. As we map phase of a cortical region directly to the onset of the presented sound, any variability will reduce the accuracy of the phase estimation in relation with the behavioral outcome. It can be expected that an area high up the processing hierarchy such as IFG has relatively high temporal variability of the neural response to speech.

To conclude, we show that word categorization depends on the oscillatory phase of the neural populations in regions where lexical and phonological information effects are typically observed. The categorization bias is mediated by the trait or linguistic unit frequency that is putatively represented in the region, perhaps through variation in sensitivity of diverse neural populations. We find a double dissociation in which the phase in STG biases participants to the low or high frequent consonant percept, while the phase in MTG biases participants to the low or high frequent word percept. These results demonstrate that oscillations provide a temporal ordering of neural activation based on the excitability of neural populations. Moreover, our study highlights the role of low frequency oscillations to organize neural activation patterns along a gradient of event probability and provide an outlook for further investigating the fundamental mechanisms that may be expressed via population rhythmic activity.

## Online Methods

### Computational modelling

We used a modified version of the Speech Tracking in a Model Constrained Oscillatory Network (STiMCON) model^17^. In this model, a population of neural nodes is modulated by an oscillation and individual nodes are additionally modulated based on their connectivity pattern with sensory input (see formula (1). In the original model individual nodes are also modulated based on feedback). Each node is governed by a non-linear inhibition function:

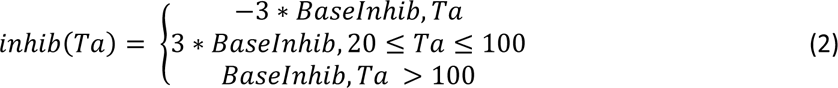

in which BaseInhib is a constant factor for the base inhibition level (set to -0.2, same as in ^17^). Initiation of the inhibition function is governed by the activation threshold (by default set on 1, but varies with neural sensitivity, see main text). First, this function creates suprathreshold activation after which nodes are inhibited. The oscillatory function is as follows:

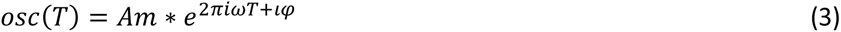

in which Am is the amplitude of the oscillator (set to 1), ω the frequency (set to 6.25Hz in accordance with ^11^), and φ the phase offset (variable). For the psychophysics experiment the phase offset is equalized with the phase of the stimulus input. In the model, sensory input is directed to two different levels of analysis, a phonetic level and a word level of analysis. Sensory input itself is modelled as a linear function ranging between 0 to 1 in arbitrary units to the individual nodes lasting half a cycle. The maximum strength of the sensory input depends on the morph level presented (see main text).

## Behavioral experiment

### Participants

In total 36 (28 female; age range = 18-40; mean age = 24.3) and 28 (20 female; age range = 20-59; mean age = 27.6) Dutch native speakers completed the session for the consonant and vowel experiment respectively. All participants reported normal hearing and did not have any history of language related disorders. Participants gave informed consent online. The study was approved by the Ethics Board of the Social Sciences Faculty of Radboud University in Nijmegen. Participants received monetary reimbursement for their participation.

### Materials

Google text-to-speech was used to utter the Dutch word *daad* (IPA [international phonetic alphabet]: /dat/; translation: *deed*) and *gaat* (IPA: /xat/; translation: *go*). We spliced the audio-file to only contain the /da/ and /xa/ parts. We max-normalized these spliced audio fragments. In praat^58^ we equalized the pitch contours of /da/ and /xa/ to lie in the middle of the original pitch contours of the two sounds. For the consonant manipulation we morphed the two sounds together by taking a weighted average of the two audio fragments in 11 spaced steps. Note that this step is different than in our original study^11^ in which we changed the formants directly, but it was necessary as we could not achieve a guttural /x/ made in Dutch by using a formant change only. While for syllable perception this procedure is not a problem (even though the sound is then closer to a /ga/ than a /xa/), for the chosen words a guttural /x/ is needed to understand the words. For the vowel manipulation, we subsequently changed the temporal modulation (for all 11 morphs) of the original /a/ sound to be 0.75 of the original duration using PSOLA^59^ (which can maintain pitch while changing temporal rate). Then we changed the spectral content of the second formant in 11 steps from 1300-1700 Hz during the vowel utterance using the burgs LPC method^60^. This morph generates the phoneme /ɑ/. We again max-normalized the output of these morphs. We spliced from the original /dat/ sound fragment the /t/, shortened it to 0.9 of the original length (to improve the sound audibility) and concatenated the /t/ at the end of all created morphs. The amplitude of the last 0.2 seconds was linearly dampened. This whole procedure created a total of 11×11 morphs, morphing between the four extreme sounds /xat/, /dat/, /xɑt/, and /dɑt/. Note that from all these morphs we only used a total of 11×4–4 sounds, that is, the sounds at which either the consonant or vowel was at its most extreme value. We choose these words as they are dissociable in vowel, consonant and word frequency (see table 1).

For the psychophysics experiment, we presented a rhythmic sequence of broadband noise at 6.25 Hz before the word. Broadband noise consisted of 50 ms (with 5 ms linear amplitude ramp up and down) between 1.1 and 3.1 kHz. Sequences lasted randomly 2, 3, or 4 seconds. Stimulus onset asynchrony (SOAs) of the word relative to the final noise in the sequence was set to be between 0.1 to 0.42 in 12 equidistant steps (covering two cycles of 6.25 Hz).

### Procedure

All procedures were done online using Django web development, running under Apache. In the first part of the experiment, we determined the most ambiguous stimulus as the morph for which participants heard on of the two extremes of the morph 50% of the time. In both experiments, this entails two morph spectra: for the consonant experiment the /dat/-/xat/ and /dɑt/-/xɑt/ spectra; for the vowel experiment the /dat/-/dɑt/ and /xat/-/xɑt/ spectra. To do so, we presented all 22 morphed sounds for the respective experiment and participants had to indicate what word they heard. A trial consisted of a silent period of 0.5 seconds followed by the presentation of the audio fragment. 0.25 seconds after the sound, participants viewed the response options and could indicate via a button press which sound they heard. Participants received two response options corresponding to the two extremes of the spectrum to which the sound morph belonged to. In total, each sound was presented 12 times, corresponding to 264 trials divided into two blocks. In total, the first part lasted about 8 minutes. Immediately after this part for both spectra a psychometric logit function was fitted, and the most ambiguous sound was determined.

The second, main, part of the experiment consisted of 14 blocks in which we presented the rhythmic sequences with the final words. In total we presented 648 experimental trials: nine repetitions, three sequence lengths, twelve SOAs, and two sound types. We added 5% of filler trials consisting of the extreme sound types (at random sequence length and SOAs) resulting in a total of 680 trials. These trials were added to test that participants were performing the task and not randomly pressing buttons and to make sure that in some instances there was also a clear correct answer. Again, after the button press there was an interval of 0.5 seconds. Throughout the experiment we adapted the ambiguous sound when participants heard the same sound too often in a row. Specifically, if participants categorized the ambiguous sound as the same word for 10 times in a row (for each spectra), we adjusted the ambiguous sound one morph step away from the perceived word.

### Behavioral analysis

For the first part of the experiment, we fitted a psychometric curve using the curve_fit function in the scipy toolbox in python and extracted the most ambiguous stimulus for each participant. We had difficulty to ensure that participants maintained an ambiguous percept either during the psychometric determination or during the main experiment. We therefore had to exclude quite a few participants from the analysis. This is likely due to the online procedure that needed to be done during the COVID pandemic to which we had to rely on the audio of the participants at their home situation. As the experimental results hinge on having an ambiguous percept, we needed to exclude participants who could not maintain an ambiguous word perception throughout the experiment. During the first part of the experiment, we could not fit an ambiguous sound that stood in between the 10^th^ and 90^th^ percentile of the morph spectrum for 7 and 4 participants for consonant and vowel experiment, respectively. Also, during the main experiment, 6 and 11 participants reported low perceived differences between the two unambiguous words in the corresponding spectrum, respectively (under 20% difference between the answers to the two unambiguous words; likely due to a failure to comply with the task or difficulties with the task itself). An additional 5 and 1 participants reached morphs outside the 10^th^ and 90^th^ percentile ranges for more than half the duration of the main experiment, respectively. This ended us with 18 and 12 participants for the consonant and vowel experiment, respectively. Trials in which morph estimations were outside of the 10^th^ and 90^th^ percentile ranges were excluded from the analysis.

For those participants who met our criteria, we created a time course across the twelve SOAs used. For each participant we subtracted the mean of the time course. Then we averaged all time-courses and fitted a 6.25 Hz sinusoid to the data (with varying amplitude and phase) and extracted the explained variance. We generated a null distribution by randomly permuting (n = 10,000) the average time course and refitting the sinusoid. One-sided p-values were extracted by comparing the proportion of the observed explained variance with the explained variance of the null distribution.

## MEG experiment

### Participants

23 Dutch native speakers (13 females; age range: 18-59; mean age = 34.3; one author participated as well) participated in the study. 22 were right-handed (one reported no preference in hand). All reported normal hearing, had normal or corrected-to-normal vision, and did not have any history of dyslexia or other language-related disorders. Participants performed a screening for their eligibility in the MEG and MRI and gave written informed consent. The study was approved by the ethical Commission for human research Arnhem/Nijmegen (project number CMO2014/288). All participants were reimbursed for their participation. One participant was excluded for not maintaining an ambiguous percept throughout the experiment.

### Procedure

Just as the behavioral experiment, the MEG experiment consisted of two parts. In the first part, we repeated the presentation of the psychometric function to determine the most ambiguous sounds. After the sound was finished the response options were immediately shown. The next sound was presented at random interval between 0.5-1.5 seconds after the response. In the MEG experiment, we only used the consonant morphs. During the main part of the experiment, we presented 50% of the time ambiguous (n = 160 per spectrum) and 50% of the time non-ambiguous sounds (n = 80 for each extreme per spectrum). All trial types were presented in pseudo-random order. After sound presentation, there was a 1 second interval before the sound options were shown. The next sound was presented at a random interval between 2 and 4 seconds after the response. After 80 sounds participants had a break. The response options of participants (pressing left or right for /d/ vs /x/, respectively) were switched halfway through the experiment to ensure that the effects were not due to motor plans. Half of the participants started with /d/ as the left option and the other half with /x/ as the left option. During the experiment, the morph was changed if the participant reported the same percept for the ambiguous sound within one spectrum for 10 consecutive answers. At the end of the experiment, we collected an auditory localizer (data not analyzed here) and a scalp digitization using the Polhemus Fastrak digitizer. All stimulus presentation was programmed in Psychtoolbox^61^ and run in the linux environment.

### MEG pre-processing

Surface-based source models from the MRI were made using grid points that were defined on the cortical sheet of the automatic segmentation of freesurfer6.0 [50] in combination with pre-processing tools from the HCP workbench1.3.2 [51] to down-sample the mesh to 4k vertices per hemisphere. The MRI was co-registered to the MEG by using the previously defined fiducials as well as an automatic alignment of the MRI to the Polhemus headshape using the Fieldtrip20211102 software [52]. Head models were based on the SPM segmentation incorporated in Fieldtrip. Regions of interest (ROIs) were the superior temporal gyrus (STG), the middle temporal gyrus (MTG) and the inferior frontal cortex (IFG). Parcellations were based on the Freesurfer parcellations.

Preprocessing of MEG involved epoching the data both between -2 and 0 seconds and -1 to +1 seconds relative to sound onset. This separate epoching was necessary to ensure that for the pre-stimulus analyses no data from the post-stimulus interval could leak into the pre-stimulus interval due to filtering during preprocessing. Data was then padded for 0.5 seconds at the end of the epoch with the last value of the epoch. Data was low-passed at 100 Hz and DFT notch filters were applied at 50, 100 and 150 Hz. Then, data was down-sampled to 300 Hz and the padded interval was removed again. ICA was performed to remove heartbeat-related signals and eye blinks and movements. On average 4.3 components (range: 3-6 components) were removed from the analysis. After that, trials with excessive noise were removed via visual inspection with an average of 12.7 removed trials (range: 4-22 trials). We calculated a common spatial filter using lcmv filter based on the post-stimulus data with a lambda of 5%. Many spatial filters have a center of the head bias, resulting in stronger activity in the center compared to the cortical surface^29^. This bias is often counteracted by having a clear baseline period for each trial to which the data is referenced to. However, for our analysis no clear baseline period can be defined as we were interested in the pre-stimulus period. Therefore, to counteract the center of head bias we used an array-gain beamformer which normalized the spatial filter^29^. This filter was applied to all single trial estimates of the pre-stimulus data. To extract a single time course representative of our ROIs we extracted the first PCA for each ROI.

### MEG analysis

We performed a time-frequency analysis on all source trials using a wavelet approach extracting frequencies from 1 to 15 Hz in steps of 0.5 Hz at widths matching 700 milliseconds for the timepoints -0.5 up to 0 seconds in steps of 0.05 seconds extracting the phase of the complex Fourier spectra. Data corresponding to the ambiguous sounds were then split according to the response of the participants based on two different contrast binning:

1) Consonant frequency binning: responses where the ambiguous word was interpreted as a word with a low frequency consonant (/xɑt/ and /xat/) versus a word with a high frequency consonant (/dɑt/ and /dat/).
2) Word frequency binning: responses where the ambiguous word was interpreted as a low frequency word (/dat/ and /xɑt/) versus a high frequency word (/dɑt/ and /xat/).

For these two contrasts, low frequency traits were labeled with a zero and high frequency traits with a one. A circular-linear correlation was performed between the pre-stimulus phase and the response of the participant for all time and frequency points. This analysis produced a correlation value and a p-value for each participant. As the exact correlation value is influenced by the number of trials included in the correlation, we performed statistics using the inverse of the normative cumulative distribution based on the p-value of the correlation of the individual participants (also see^62^). Group statistics were performed by statistically testing these z-values against zero using a one-sample t-test. To control for multiple comparisons across all time-frequency points we performed cluster-based permutation tests^31^.

To further inspect the effect of the two contrasts we split the data based on all four possible response options. Then, we extracted for each participant and for each of the possible perceived words the average phase at which participants reported perceiving that specific word. We calculated the phase difference between the average phase at which participants perceived the word /dɑt/ and the other three options. The logic of this analysis was as follows: /dɑt/ has high frequency features for all investigated feature dimensions. Words that also have a high frequency content should therefore show a phase difference of zero with /dɑt/, but words that have a low frequency content should show a phase difference of π with /dɑt/. We statistically tested whether the phases were non-uniform around the expected phase using the v-test statistic^63, 64^. This test will show a significant effect only when the data is both non-uniform and the phase is around the expected phase.

For the power analysis, we used the same wavelet approach as for the time-frequency plots (Figure 4C) and converted the data in z-scores across the whole time-frequency window. For the power spectra (Figure 4B), we cut the data from -1 to 0, padded the data out to 5 seconds and extracted the power using Hanning tapers.

## Acknowledgements

AEM was supported by the Lise Meitner Research Group “Language and Computation in Neural Systems”, by NWO Vidi grant 016.Vidi.188.029, and by the Language in Interaction Consortium which is funded by NWO Gravitation Grant 024.001.006.

