## Supplementary Figure for "Phase-dependent word perception emerges from region-specific sensitivity to the statistics of language"

### Supplementary figures

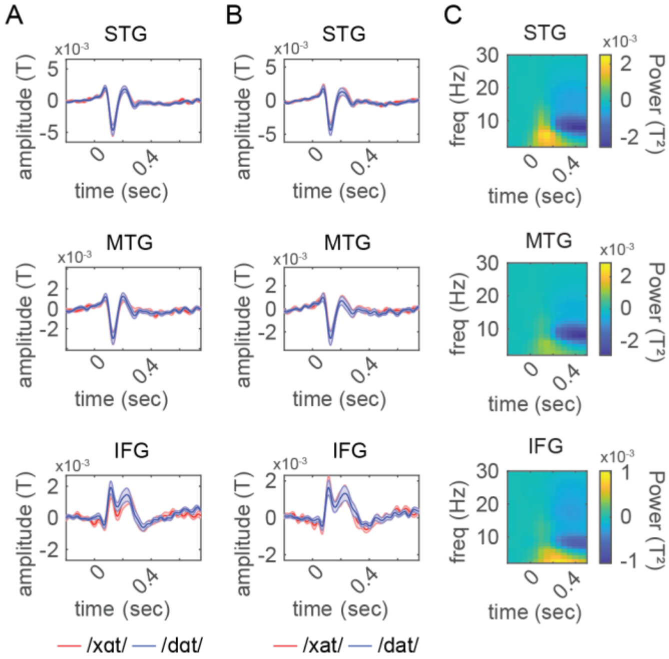


**Supplementary figure 1.** Responses in regions of interest. A) Response to ambiguous stimulus /xɑt/-/dɑt/. B) Response to ambiguous stimulus /xat/-/dat/. C) Time-frequency response averaged across all stimuli. STG = superior temporal gyrus. MTG = medial temporal gyrus. IFG = inferior frontal gyrus.
